# Guess till correct: Gungnir codec enabling high error-tolerance and low-redundancy DNA storage through substantial computing power

**DOI:** 10.1101/2025.08.29.673174

**Authors:** Jingcheng Zhang, Lei Chen, Jinlin Sun, Shumin Li, Yekai Zhou, Zhenqin Wu, Can Li, Zhenxian Zheng, Ruibang Luo

## Abstract

DNA has emerged as a compelling archival storage medium, offering unprecedented information density and millennia-scale durability. Despite its promise, DNA-based data storage faces critical challenges due to error-prone processes during DNA synthesis, storage, and sequencing. In this study, we introduce Gungnir, a codec system using the proof-of-work idea to address substitution, insertion, and deletion errors in a sequence. With a hash signature for each data fragment, Gungnir corrects the errors by testing the educated guesses until the hash signature is matched. For practicality, especially when sequenced with nanopore long-read, Gungnir also considers biochemical constraints including GC-content, homopolymers, and error-prone motifs during encoding. *In silico* benchmarking demonstrates its outperforming error resilience capacity against the state-of-art methods and achieving complete binary data recovery from a single sequence copy containing 20% erroneous bases. Gungnir requires neither keeping many redundant sequence copies to address storage degradation, nor high-coverage sequencing to address sequencing error, reducing the overall cost of using DNA for storage.

## Introduction

DNA is an attractive storage medium with great potential to meet the growing demand for archiving. In DNA archiving, binary information is encoded into sequences composed of A/C/G/T nucleotides according to a defined protocol. These sequences are then synthesized into fragments and stored *in vitro*. In contrast to conventional information storage media, DNA-based storage exhibits an ultra-high storage density (∼6 orders of magnitude greater than traditional media ^1^) and remarkable durability (maintaining molecular stability for millennia ^2^ under proper storage conditions). Additionally, DNA can be passively preserved without additional energy consumption. These unique properties make DNA a compelling solution for next-generation archival storage systems.

Despite its advantages, DNA archiving poses great challenges due to the errors introduced throughout its lifecycle, including synthesis, degradation, and sequencing. During DNA synthesis, multiple DNA sequences (oligos) of 100–300 nucleotides ^3^ are produced for each encoded sequence, and might introduce 0.5–2% synthetic errors ^4^. However, these errors might be amplified during polymerase chain reaction (PCR) ^4^ and accumulate over time due to molecular degradation ^5^. Sequencing technologies, such as short-read ^6^ and long-read ^7^ sequencing, which are employed to recover the encoded data, can also introduce additional errors. Consequently, a more robust codec system is necessary ^8^ to correct the common 3– 5% erroneous bases.

In particular, three types of errors might corrupt the encoded information, including substitutions (replacing bases), insertions (adding bases), and deletions (removing bases). To fix these errors, existing codec systems used decoding strategies that convert insertions and deletions (Indels) to substitutions ^9, 10^ or sequence losses ^11, 12, 13, 14, 15^, and used computer erasure codes, such as DNA fountain ^16^ and Reed-Solomon (RS) codes, to reconstruct the original information. These existing systems cannot effectively address Indels. To compensate, they require excessive physical redundancy (5–10× DNA copies, 50–100× high-coverage sequencing ^17^) or logical redundancy (an information density of less than 1 bit per base) to address these errors. Moreover, experimental results show that existing codec systems fail when the overall error rate exceeds 5.25% ^8^—a threshold frequently exceeded in the long lifecycle ^18^ of DNA storage.

Therefore, to enable efficient, durable and reliable DNA-based long-term storage, codec systems with high error tolerance that can accurately correct substitutions and Indels are essential. To address these challenges, we developed a novel approach drawn from the blockchain proof-of-work concept ^19^. In blockchain systems, miners repeatedly guess combinations of transactions and nonces to find one that produces a valid hash signature, thereby earning rewards such as Bitcoin ^20^. Similarly, in a DNA-based codec system, a decoder can iteratively generate candidate sequences that follow predefined character generation rules until the resulting hash signature matches the target. This cryptographic framework ensures correct sequence recovery given substantial computing power, while the inherent difficulty of cryptographic hashing prevents false positives.

In this work, we proposed Gungnir, a DNA-based storage codec system named after the weapon of the Norse god Odin, known for its unwavering accuracy. Gungnir substantially raises the upper limit of the maximum tolerable error rate of a codec system—even information stored in sequences with 20% erroneous bases (equivalent to 406 years of degradation at 13 degrees Celsius ^21^) can be fully recovered. To our best knowledge, Gungnir is the first codec system that can effectively deal with excessive Indels in a sequence, which is common in DNA-based storage. Gungnir supports adjustable configurations to maintain a balance between information density and error tolerance.

Compared to existing methods, Gungnir achieves the same performance while requiring only half the DNA bases. Additionally, Gungnir can employ base preference tables for generated sequences; for instance, it can produce sequences with a 28.4% reduction in expected Nanopore sequencing errors, albeit at the cost of some acceptable error tolerance capacity. The outstanding results suggest that Gungnir has great applicability in diverse DNA-based storage scenarios, including both long-term archiving and high-frequency retrieval scenarios. With affordable computational resources, Gungnir facilitates robust information recovery while maintaining low redundancy, making it a reliable solution for DNA-based data storage codec systems.

## Results

### Overview of Gungnir Codec System

**Figure 1a** illustrates the workflow of DNA-based digital information storage, which comprises two key stages: encoding and decoding. Encoding involves transcoding binary fragments into DNA sequences, and decoding is to reconstruct the original information from sequences with errors. When encoding a file, such as an image, video, or audio, the encoder is designed to transcode the binary stream into DNA sequences using a minimal number of bases. The decoder, on the other hand, is designed to correctly reconstruct data from a sequence with a number of erroneous bases. However, as the error rate increases, the codec system may fail to maintain a balance between information density and error tolerance. This is because the vast array of potential correct sequences becomes more challenging to distinguish, necessitating excessive redundancy for error correction. This redundancy ultimately leads to suboptimal information density ^22^.

**Figure 1.**
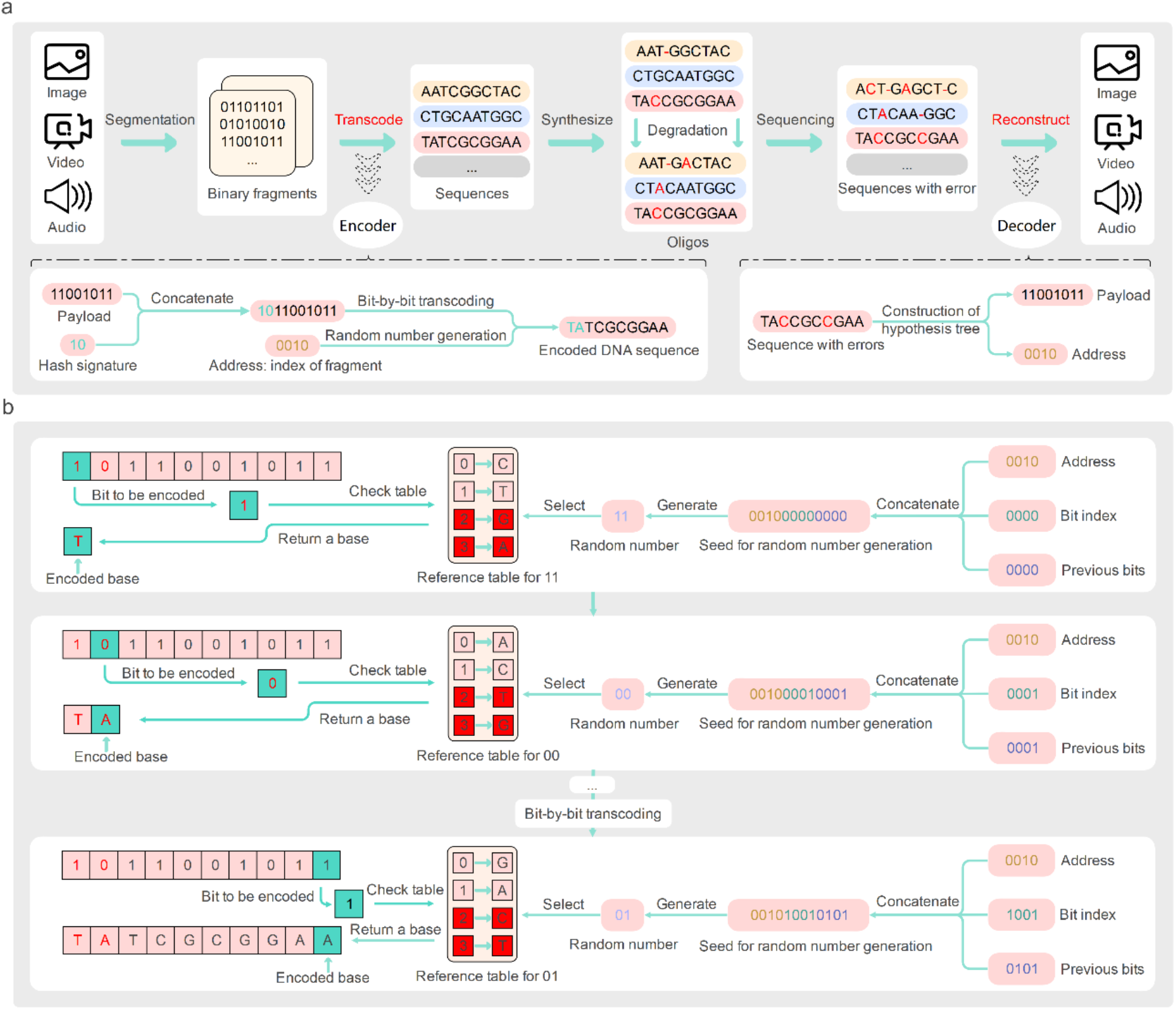
Gungnir codec overview and its encoding algorithm. a. General workflow of the Gungnir codec. First, data is segmented into fragments of binary (also referred to as payloads). The encoder then transcodes the concatenation of each fragment and its hash signature into a nucleotide sequence of length 100 to 300. The sequences are then synthesized into oligos. Errors are then introduced and accumulated during synthesis and storage. When data retrieval is required, the oligos are sequenced and reconstructed by the decoder. b. Details of the Gungnir encoder. In the bits to be encoded, the bits of hash signature are labeled in red, and the bits of payload are labeled in black. The encoder transcodes bits into nucleotides bit-by-bit, with which nucleotide to be used for a bit decided by one of the preset reference tables, chosen by a random number generated from a seed constructed as the concatenation of address, bit index, and previous bits.

This observation forms the core principle of Gungnir: if a decoder can produce a decoded binary fragment from a sequence, it should be mathematically proven to be reliable and accurate. Guided by this principle, Gungnir introduces a hash signature for each data fragment, serving as a proof of identity, as detailed in the **Methods** section **‘Gungnir Transcoding’**. As shown in **Figure 1a**, the Gungnir encoder transcodes the hash signature of each data fragment into nucleotides, along with the data bits (payload). During decoding, the Gungnir decoder exhausts all possible payloads and hash signatures that correspond to a sequence with errors (see **Figure 2a**) within a specified *W*_*max*_ (i.e., maximum hypotheses allowed). Only hypotheses that successfully pass the hash signature validation are outputted. If no valid results are generated, the Gungnir decoder increases the *W*_*max*_ and repeats the decoding algorithm until obtaining correct hypothesis. In this manner, when sufficient bits for the hash signature and adequate computing power for the proof-of-work architecture are available, the Gungnir decoder can reliably identify the correct binary stream as output.

**Figure 2.**
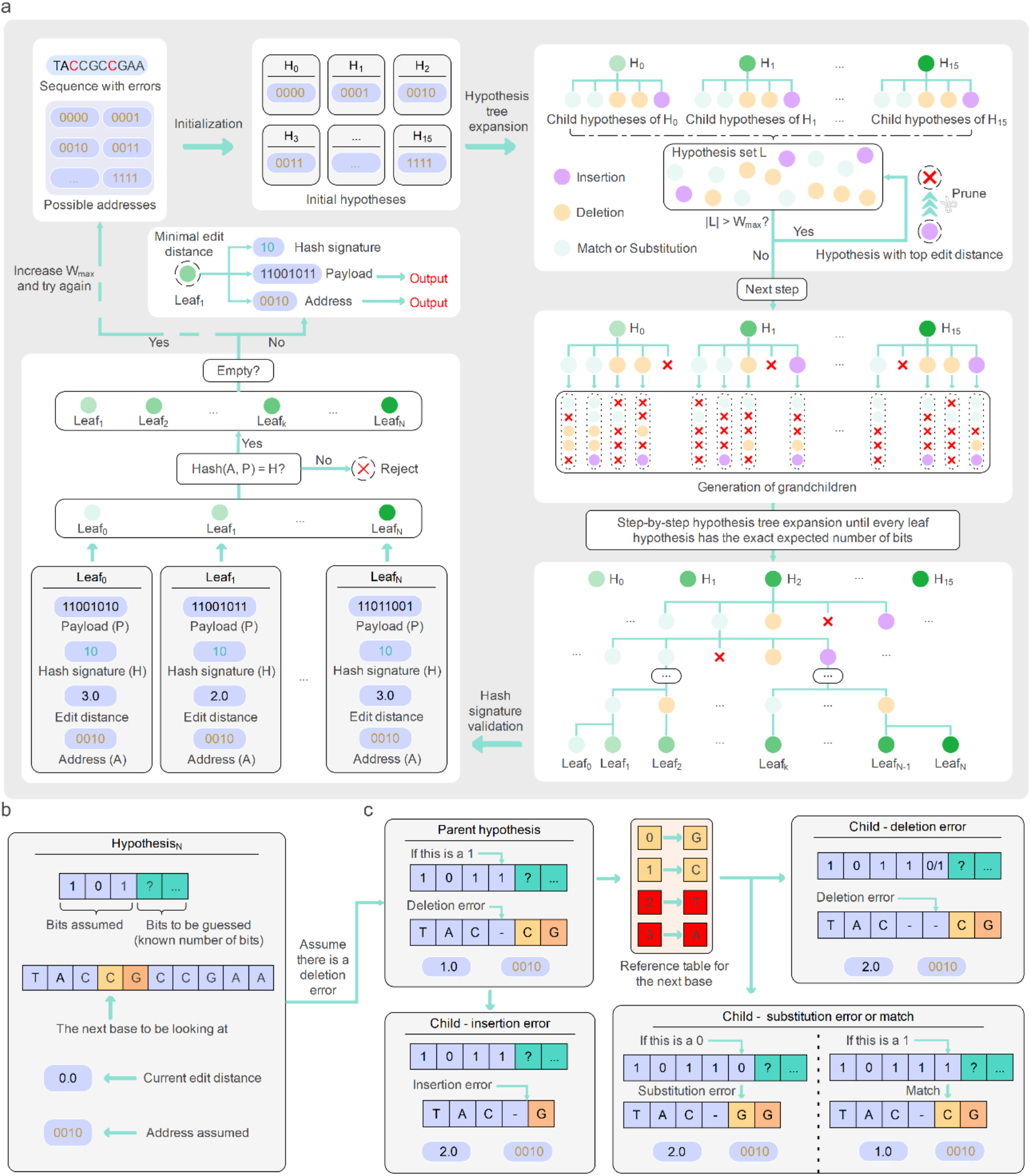
Details of the Gungnir decoder. a. The decoding pipeline. Firstly, the Gungnir decoder assigns an initial hypothesis for all remaining possible addresses (those addresses with successfully decoded payload are ignored). In the subsequent iterations, every hypothesis at a layer generates five more children hypotheses in the next layer. When the number of hypotheses at a layer *i* exceeds a configurable *W*_*max*_ (maximum hypotheses allowed), the decoder prunes hypotheses with the highest edit distance (between the hypothesis and the original input) in the tree until *i* falls below *W*_*max*_. Once all leaves in the tree have the exact expected number of bits (determined in encoding and same for all payloads), the decoder performs hash validation. Wrong hypotheses failing hash validation are pruned. Hash collision is rare but happens when the configurable hash signature size is set to small, or the sequence error rate is high. The decoder outputs the candidate that passes hash validation with the minimal edit distance as the decoding result. If no decoding result is generated, the decoder increases *W*_*max*_ and try again. b. Components of a hypothesis. c. How five children hypotheses are generated from a parent hypothesis, with five possible assumptions of what happened to the sequence at a position, including 1) a match, 2) a substitution, 3) an inserted base, 4) a deleted base with a guess of the next bit to be 0, and 5) a deleted base with a guess of the next bit to be 1.

### Performance of the Gungnir Codec System

#### Error Tolerance of Gungnir in Single-read DNA Data Reconstruction

The robustness of a DNA-based codec system is primarily determined by its ability to correct errors. To evaluate the data recovery performance of the Gungnir codec system, we compared it to five state-of-the-art codec systems: Church ^11^, Grass ^13^, Yin-yang ^14^, DNA-Aeon ^15^, and HEDGES ^9^. All systems were configured with consistent constraints: a maximum homopolymer length of 3 and a GC content between 40–60%.

To assess each codec’s native error tolerance capacity with single-read coverage, we conducted tests using single-molecule reads without consensus correction and established two error profiles for evaluation. In error profile 1, we maintained equal proportions of substitutions, insertions, and deletions errors and increased the errors to each sequence until the overall error rate reached 20%. This *in silico* benchmark aimed to evaluate the error tolerance capabilities of the different codec systems. In error profile 2, we fixed a specific Indel rate (0.54% deletions and 0.23% insertions, based on the *in vitro* test data from HEDGES) and incrementally added substitutions to each sequence until the error rate reached 20%. This error profile mimicked real-world conditions where storage-induced degradation primarily generates substitutions ^4^.

**Figure 3a** illustrates data recovery performance when with the same proportions of substitutions, insertions, and deletions errors (error profile 1), with parameters and settings detailed in **Supplementary Text 1**. The Church, Grass, and Yin-yang codec systems were optimized for substitutions, and they failed to achieve 100% data recovery when insertions or deletions were introduced. Notably, when the total error rate reached 1%, their data recovery rates fell between 35.3% and 48.0%. This trend persisted as the error rate increased to 3%. DNA-Aeon, in collaboration with its outer DNA fountain ^16^ code, maintained perfect data recovery until 3.9% error rate and failed completely by 4.2% error rate. Similarly, HEDGES, in collaboration with its outer Reed-Solomon code, achieved 100% data recovery until an error rate of 5.25%, after which its performance deteriorated significantly around 6% to 7.5% error rate. In contrast, Gungnir maintained 100% data recovery even at 20% error rates, demonstrating superior robustness against Indels and representing a 3–5× improvement over existing systems.

**Figure 3.**
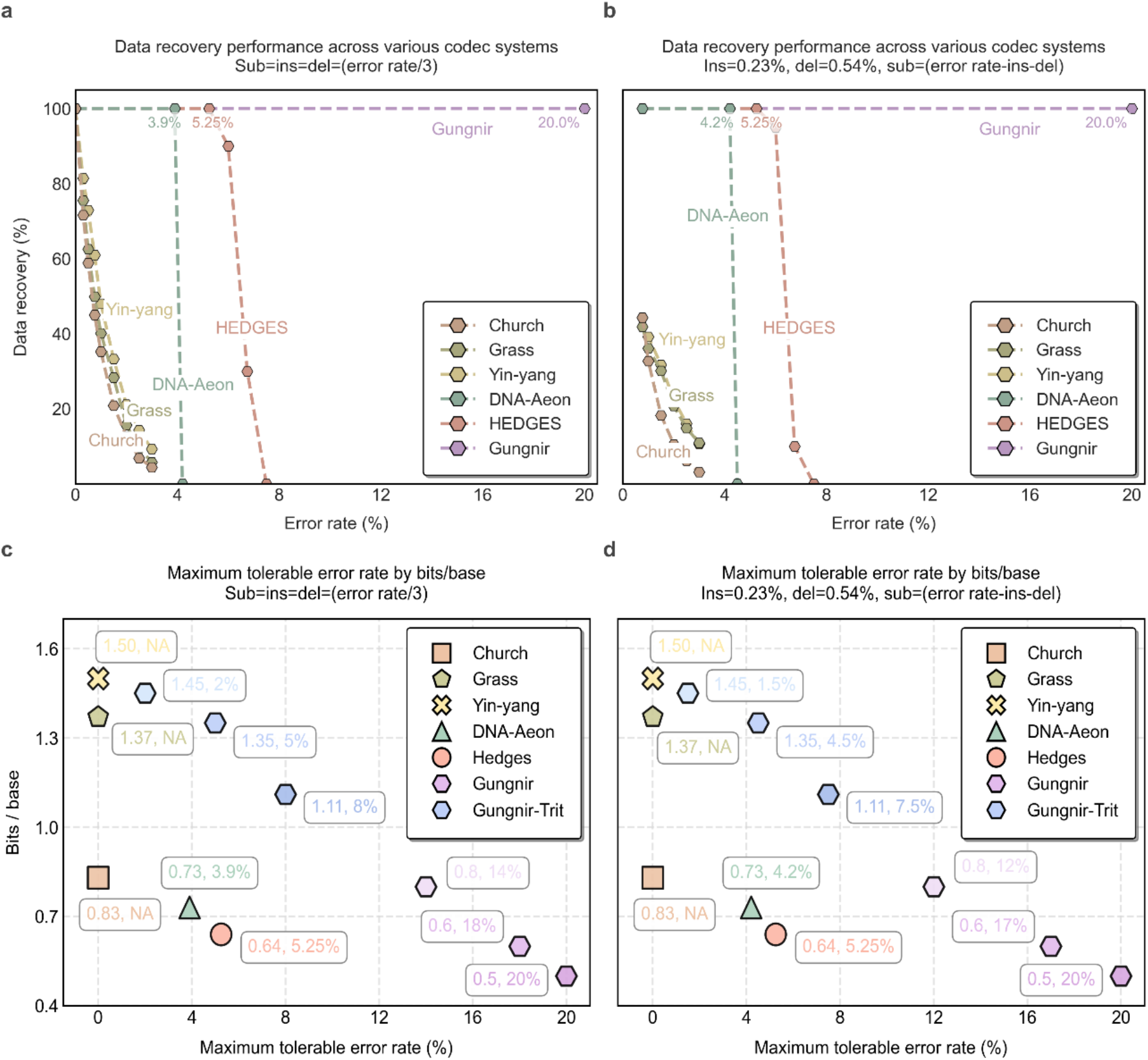
Error tolerance performance of Gungnir. a-b. A comparison of the percentage of payloads fully recovered at different error rates up to 20% using different codecs. All codecs used default settings. The default bits/base of Gungnir is 0.5. c-d. A comparison of the maximum tolerable error rate at different bits/base of different codecs, when requiring payloads to be fully recovered. For both a and b, two different error profiles were tested. Sub, ins, and del stand for substitution, insertion, and deletion, respectively. See **Supplementary Table 1** and **Supplementary Table 2 for details**.

**Figure 3b** illustrates the data recovery performance of different codec systems when fixing insertion and deletion rates at 0.23% and 0.54% respectively (error profile 2). When no substitutions were introduced, Church, Grass, and Yin-yang recovered 41.9% to 44.3% of the data. However, their data recovery rates decreased significantly when the number of substitutions increased, falling to less than 11% when with 2.23% substitutions (3% total errors). Like observations from error profile 1, DNA-Aeon maintained perfect data recovery until 3.43% (4.2% total errors) substitution rate and then dropped sharply to zero. HEDGES maintained 100% data recovery up to a substitution rate of 4.48% (5.25% total errors). In contrast, Gungnir achieved 100% recovery even with up to 19.23% substitutions (20% total errors). This exceptional performance demonstrates Gungnir’s capacity to recover information from degraded DNA samples in archival storage applications.

### Benchmarking Information Density in DNA Storage Codec

Beyond error tolerance capacity, it is also crucial to optimize information density for DNA-based codecs. The primary measure of information density is the number of bits that each base can encode (bits/base). Noteworthy, the metric should account for all redundancy, including both inner code overhead and supplementary sequences from outer error correction code.

In **Figure 3c and 3d**, we analyzed the correlation between the maximum tolerable error rate and information density for all benchmarked codec systems. Considering the redundancy introduced by indexing and error correction, the actual information densities of the benchmarked codec systems ^9, 11, 13, 14, 15^ range from 0.64 to 1.50 bits/base. For the state-of-the-art codec system HEDGES, the actual information density is ∼0.64 bits/base, with a maximum tolerable error rate of 5.25% in both error profiles. In contrast, Gungnir tolerated 5% and 4.5% errors in an information density of 1.35 bits/base in error profile 1 and 2, respectively, achieving ∼2 times the information density of HEDGES while maintaining similar levels of error tolerance. At 0.6 bits/base, Gungnir tolerated 18% and 17% errors in error profile 1 and 2, surpassing HEDGES’ tolerance by more than three times.

### Computing Resources Analysis with Different Error Rates

In the simulation presented in previous sections, we deliberately excluded computational constraints to facilitate exhaustive hypothesis testing. This approach enabled the decoder to iteratively increase the *W*_*max*_ (maximum hypotheses allowed) during exhaustive searching, as detailed in the **Methods** section **‘Decoding Algorithm of Gungnir’**. However, for practical deployment, it is essential to comprehend how Gungnir manages memory usage and processing time during iterative decoding. In this section, we encoded a 19.4 KB text file containing the fairy tale “The Ugly Duckling” into 3,108 DNA sequences with 100 bases, achieving an information density of 0.5 bits/base. In accordance with error profile 1, we maintained equal proportions of substitutions, insertions, and deletions errors while gradually increasing the total error rates from 10% to 20%, thereby focusing on the changes in computational resources required for decoding. These experiments were conducted on a dual 32-core Xeon Platinum 8369C server equipped with a maximum of 512GB of RAM.

For each simulation, we began with a smaller *W*_*max*_ value of 100k, which required ∼100MB memory. If a valid guess was not found, larger *W*_*max*_ settings (1M, 5M, 20M, and 200M) were employed in subsequent rounds (up to five rounds). The maximum *W*_*max*_ required ∼200GB memory, but it was only activated when the error rate reached at least 18% (as illustrated in **Figure 4a and 4b**), which is not a frequent occurrence during DNA synthesis and sequencing. The maximum memory consumption was ∼20GB when the error rate was ⩽ 16% and only 1GB when the error rate was ⩽10%, both of which are well within the capacities of modern personal computers.

**Figure 4.**
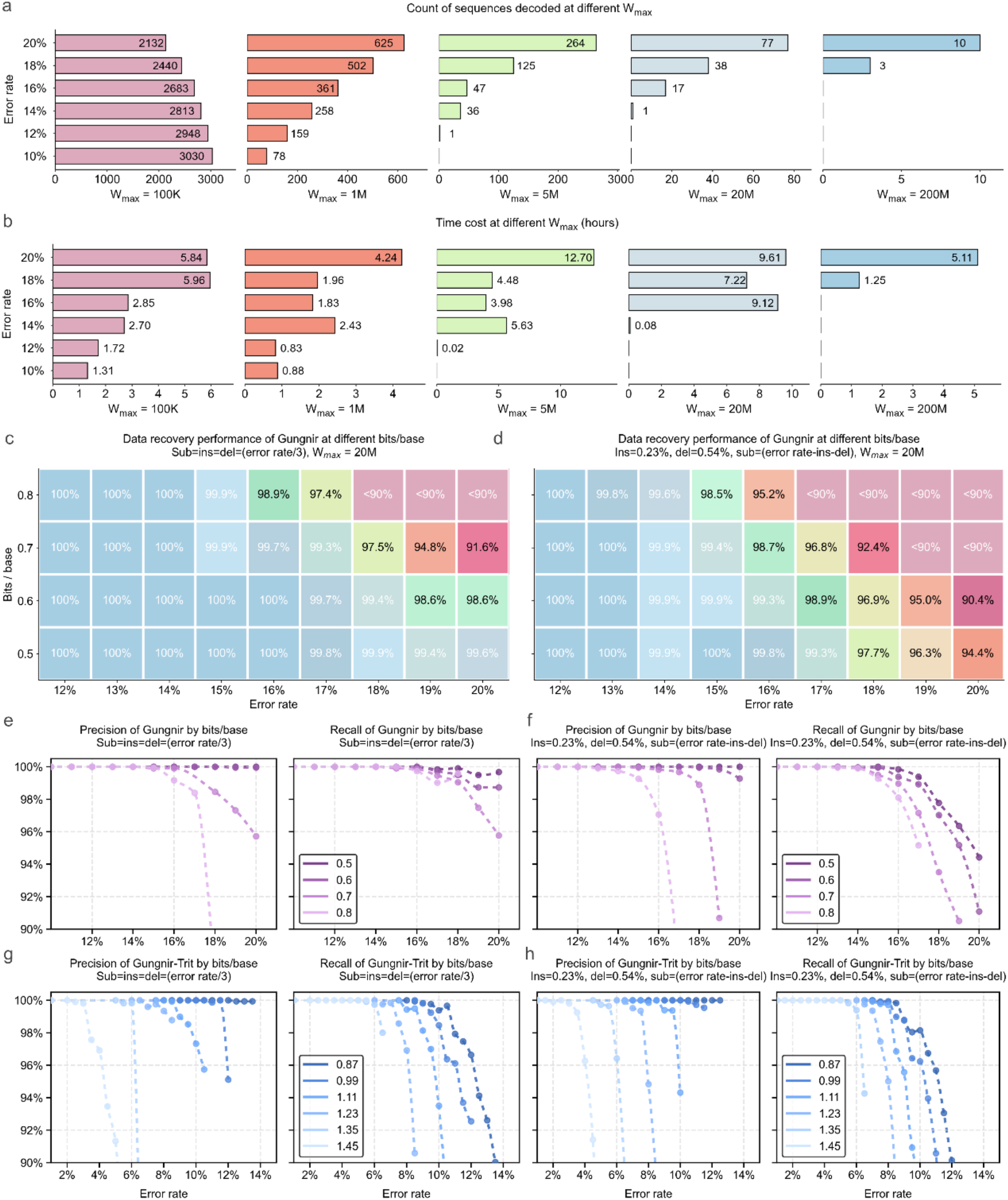
Computational cost analysis of Gungnir. a. Within total 3,108 sequences to be decoded, the figure shows the count of sequences that can be successfully decoded until which *W*_*max*_ (maximum hypotheses allowed) is set. Error rates from 10% to 20% were tested. At 10%, all sequences were successfully decoded with a *W*_*max*_ of 100k, while at 20%, although all sequences were successfully decoded, 10 of them required a *W*_*max*_ of 200M. A larger *W*_*max*_ leads to a larger memory consumption. *W*_*max*_ of 100k required ∼100MB memory, while *W*_*max*_ of 200M required ∼200GB memory. b. Time consumption of the same experiment shown in a. c-d. Data recovery performance of Gungnir at different bits/base and two different error profiles with constrained memory (*W*_*max*_=20M, ∼20GB). e-f. Precision and recall of Gungnir at different bits/base and two different error profiles with constrained memory (*W*_*max*_=20M, ∼20GB). g-h. Precision and recall of Gungnir-Trit at different bits/base and two different error profiles with constrained memory (*W*_*max*_=20M, ∼20GB). The data points of a-b and c-h are given in **Supplementary Table 3** and **Supplementary Table 4**, respectively.

In terms of the runtime, with lower error rates (⩽5%), the time consumption of Gungnir is no more than 15 minutes, which is comparable to other codecs (⩽10 minutes). When the error rate reached 10%, the time consumption increased to ∼2h. As the error rate further increased, it is more challenging to decode the sequence, requiring additional computational resource. For instance, with 14% error rate, the time consumption increased to ∼11h, with the fourth round (*W*_*max*_=20M, ∼20GB) activated. When the error rate eventually reached 20%, the total time consumption exceeded 37 hours, with over 73% of this time concentrated in the last three rounds. Following the rapid development in the computing power, we anticipate that the time required for this process will be further reduced in the near future due to advancements in hardware industry and algorithms.

### Performance Analysis of Gungnir with Limited Computing Resources

To satisfy the needs and constraints for diverse application scenarios and computing platforms, Gungnir offered various configurable options to balance information density, error tolerance, and computing resources.

### Flexible Information Density Options

DNA-based storage can represent two bits using a nucleotide base A/C/G/T, resulting in a maximum information density of 2 bits/base. However, this approach leaves no room for error correction. Gungnir addresses this limitation by using nucleotide bases to encode either payload data or hash signatures. By controlling the level of redundancy, Gungnir offers adjustable configurations for information densities, ranging from 0.5 to 1.45 bits/base. For high error rate scenarios (>10%), users can choose densities between 0.5 and 0.8 bits/base to ensure accurate reconstruction. Lower error rate applications (<10%) may utilize a higher density option to minimize the amount of DNA sequences needed for storage.

Alternatively, Gungnir-Trit, a variant that converts input from binary to trinary before transcoding, allows for even higher information densities of 0.87 to 1.45 bits/base. These different options allow users to balance storage efficiency and error tolerance based on their needs.

For fair comparison, following experiments were conducted with a fixed sequence length of 100 and the *W*_*max*_ (maximum hypotheses allowed) was set to 20M (∼20 GB memory). The file “The Ugly Duckling” was encoded into DNA sequences with 40%–60% GC content and a maximum homopolymer length of 3. We increased the error rates for two error profiles until data recovery fell below 90% or total error exceeded 20%.

**Figure 4c and 4d** demonstrate the data recovery performance of Gungnir at various bits/base levels, and **Figure 4e and 4f** illustrate the corresponding precision and recall. Sequences that are decoded but with incorrect output reduce precision, while sequences that are unable to be decoded with the given resources reduce recall. When the information density ranged from 0.5 to 0.8 bits/base, Gungnir consistently achieved 100% correct decoding in error profile 1 for error rates below 14%, and in error profile 2 with error rates below 12%. However, as the error rates increased further, data recovery gradually declined due to two main factors: decoding failures and decoding mistakes. Decoding failures result in no output, occurring when insufficient computational power is available, which subsequently leads to a decrease in recall. These failures are generally reversible, as additional computational resources can restore them. In contrast, decoding mistakes lead to incorrect outputs, primarily due to hash collisions caused by insufficient bit length in the hash signature, which reduces precision. These mistakes are typically irreversible, leading to incorrect information being decoded.

**Figure 4g and 4h** demonstrates the precision and recall of Gungnir-Trit, and the corresponding data recovery performance is presented in **Supplementary Figure 1**. In comparison with Gungnir, Gungnir-Trit’s trit-per-base encoding increases hypothesis space, making the decoder more sensitive to base errors. Despite that, when the error rate reached 5% in error profile 1 and the information density was set to 1.35 bits/base, Gungnir-Trit maintained its precision and recall at 100%. The results demonstrate Gungnir-Trit’s capacity to correct the typical 3–5% erroneous bases while preserving a better information density (1.35 bits/base vs 0.8 bits/base) than Gungnir.

### Performance at Different Sequence Length

In addition to different information density configurations, Gungnir also offers flexibility with arbitrary sequence lengths. Since common oligonucleotide lengths range from 100 to 300 nucleotides, we conducted simulations with sequence lengths of 100, 200, and 300, at error rates ranging from 5% to 20% in error profile 1. Gungnir’s information density is fixed at 0.9 bits per base, where each base represents one bit, with only 10% of the bits allocated for the hash signature. Additionally, the *W*_*max*_ was set to 100k (∼100MB memory), which is more resource-constrained compared to the experiments conducted in the previous sections to highlight the impact of sequence length on Gungnir.

The precision and recall using different sequence lengths are presented in **Figure 5a and 5b**. Notably, longer sequences demonstrate higher precision at the same error rate, indicating greater error tolerance capacity of Gungnir. For instance, as shown in **Figure 5a**, a sequence length of 300 nucleotides maintained 100% precision at a 20% error rate with a configuration of 0.9 bits/base, while a sequence length of 100 nucleotides achieved the same precision only when the error rate ⩽5%. Nevertheless, longer sequences result in lower recall, reflecting the increased computational power required for decoding. Specifically, under the same error rate, longer sequences generated more potential correct hypotheses during decoding and required more time for validation. As demonstrated in **Figure 5b**, at a 20% error rate, the recall for a sequence length of 100 nucleotides was 99.0%, while for a sequence length of 300 nucleotides, it dropped to only 3.0% due to the *W*_*max*_ constraint.

**Figure 5.**
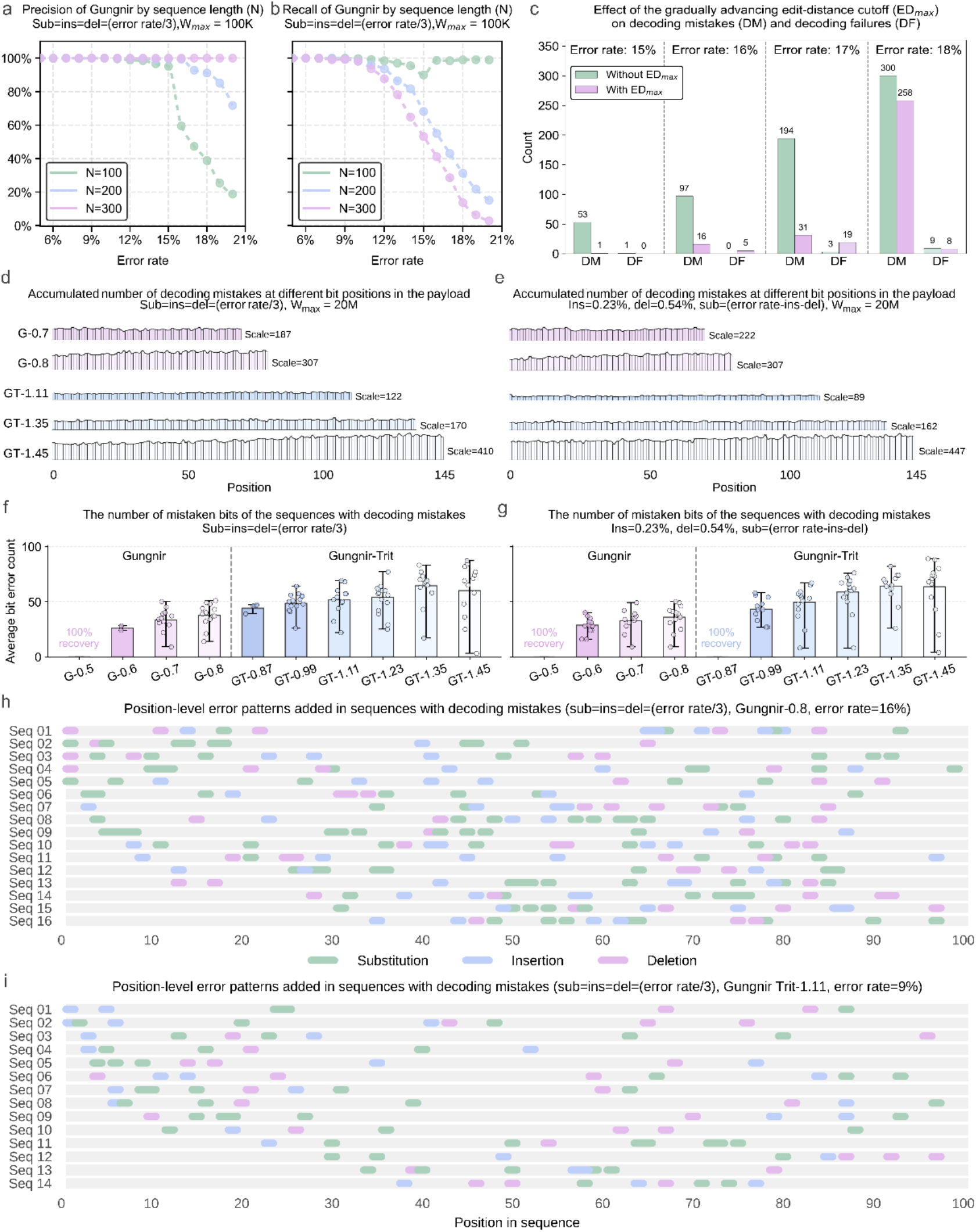
Analysis of the characteristics of decoding mistakes. Subfigures a-b tested on sequences of 100, 200, and 300 bases long, while c-i all tested on 100 bases. a-b. Precision (a) and recall (b) of Gungnir at different sequence lengths. 0.9 bits/base and *W*_*max*_=100k were used. c. A comparison of the number of decoding mistakes (DM) and decoding failures (DF) when a gradually advancing edit-distance cutoff (*ED*_*max*_) is applied or not. Gungnir 0.8 bits/base, error profile 1 (equal number of substitutions, insertions and deletions) and *W*_*max*_=20M were used. d-e. Histograms showing the accumulated number of decoding mistakes at different bit positions in the payload, using Gungnir (G) and Gungnir-Trit (GT) with varied bits/base settings. The number of bits per payload of each setting is calculated as 100 bases x bits/base, e.g., GT-1.45 has 145 bits. Two error profiles were tested. f-g. Boxplots showing the number of mistaken bits of the sequences with decoding failure. Two error profiles were tested. h-i. The exact error types and their positions among the sequences with decoding failure under two settings. The data points of a-b and c are given in **Supplementary Table 5** and **Supplementary Table 6**, respectively.

### Early Rejection of Incorrect Hypotheses Due to Hash Collision

During decoding, if the decoder outputs a valid result with abnormally high edit distance (e.g., edit distance >30 in a sequence of length 100), it is more likely to be a hash collision rather than a correct output. To reduce decoding mistakes caused by hash collision, we set an edit distance cutoff, denoted as *ED*_*max*_, and reject hypotheses with accumulated edit distance exceeding *ED*_*max*_, as detailed in the **Methods** section **‘Decoding Algorithm of Gungnir’**. For sequences that remain undecodable after multiple attempts, the decoder increases *ED*_*max*_ and restarts the decoding process. This design minimizes decoding mistakes resulting from hash collision in early attempts.

**Figure 5c** illustrates the impact of the edit distance cutoff *ED*_*max*_ on decoding mistakes and failures at varying error rates. The simulation was conducted in error profile 1, with a configuration of 0.8 bits/base and an error rate ranging from 15% to 18%. The *W*_*max*_ was set to 20M, which required ∼20 GB memory. The results showed that applying *ED*_*max*_ substantially reduced decoding mistakes, as many mistakes caused by hash collisions were effectively rejected in early attempts, which could then be corrected in subsequent attempts. This result demonstrates the necessity of edit distance cutoff for the reliability of the decoder.

### Analysis of the Characteristics of Decoding Mistakes

Decoding errors can be categorized into two types: failures and mistakes. While a decoding failure results in no output, a decoding mistake remains undetected and produces incorrect data bits (payload). In this section, we investigated instances of undetected decoding mistakes, and analyzed their underlying causes and characteristics.

### Distribution of Mistaken Bits

Existing decoding algorithms based on the hypothesis tree ^9, 10^ may encounter a challenge where the last few bits are prone to mistakes. This is because when the decoder attempts to guess these bits, there is insufficient subsequent information to effectively prune erroneous branches. In contrast, the primary cause of decoding mistakes in Gungnir shifted from untimely pruning to hash collisions. For configurations with lower information density, where more bits are allocated for hash signature, the distribution of mistaken bits became more uniform across different positions (**Figures 5d and 5e)**. Furthermore, as depicted in **Figures 5f and 5g**, the number of mistaken bits decreased proportionally to the reduction in information density. These observations align with our expectations: a longer hash signature (with its lower information density) enhances the decoder’s resilience to errors.

### Error Patterns in Sequences with Decoding Mistakes

Even when the total error rate remains constant, distributions of errors may pose challenges for the decoder. **Figure 5h and 5i** illustrate some cases where Gungnir and Gungnir-Trit yielded decoding mistakes. The distribution of errors in the sequences were plotted. Notably, in such sequences, errors were densely clustered, with every sequence exhibiting a local region of high error concentration. These dense error clusters caused the decoder to incorrectly prune correct branches when navigating these areas. Consequently, if another potential sequence that resulted from hash collision was available, it is possible to be mistakenly identified as a valid output.

### Gungnir-ONT: Frequent Data Retrieval Using Nanopore Long Reads

In addition to long-term archiving, performance for frequent data retrieval is also crucial ^2^. In this scenario, data is frequently read from oligos using sequencing technologies, in which sequencing errors can greatly increase costs. To minimize errors associated with sequencing, we proposed Gungnir-ONT, a variant that utilize a base preference table to reduce errors in Nanopore sequencing platforms (detailed in the **Methods** section **‘Character Generation Rules for Gungnir-ONT’**), and evaluated its performance.

As depicted in **Figure 6b**, the GC content of sequences generated using Gungnir, Gungnir-Trit, and Gungnir-ONT was effectively constrained within the acceptable range of 0.4 to 0.6. Notably, the majority of sequences generated by Gungnir-ONT exhibited a GC content between 0.4 and 0.45, to fit the optimal GC content range for Nanopore sequencing technology.

**Figure 6.**
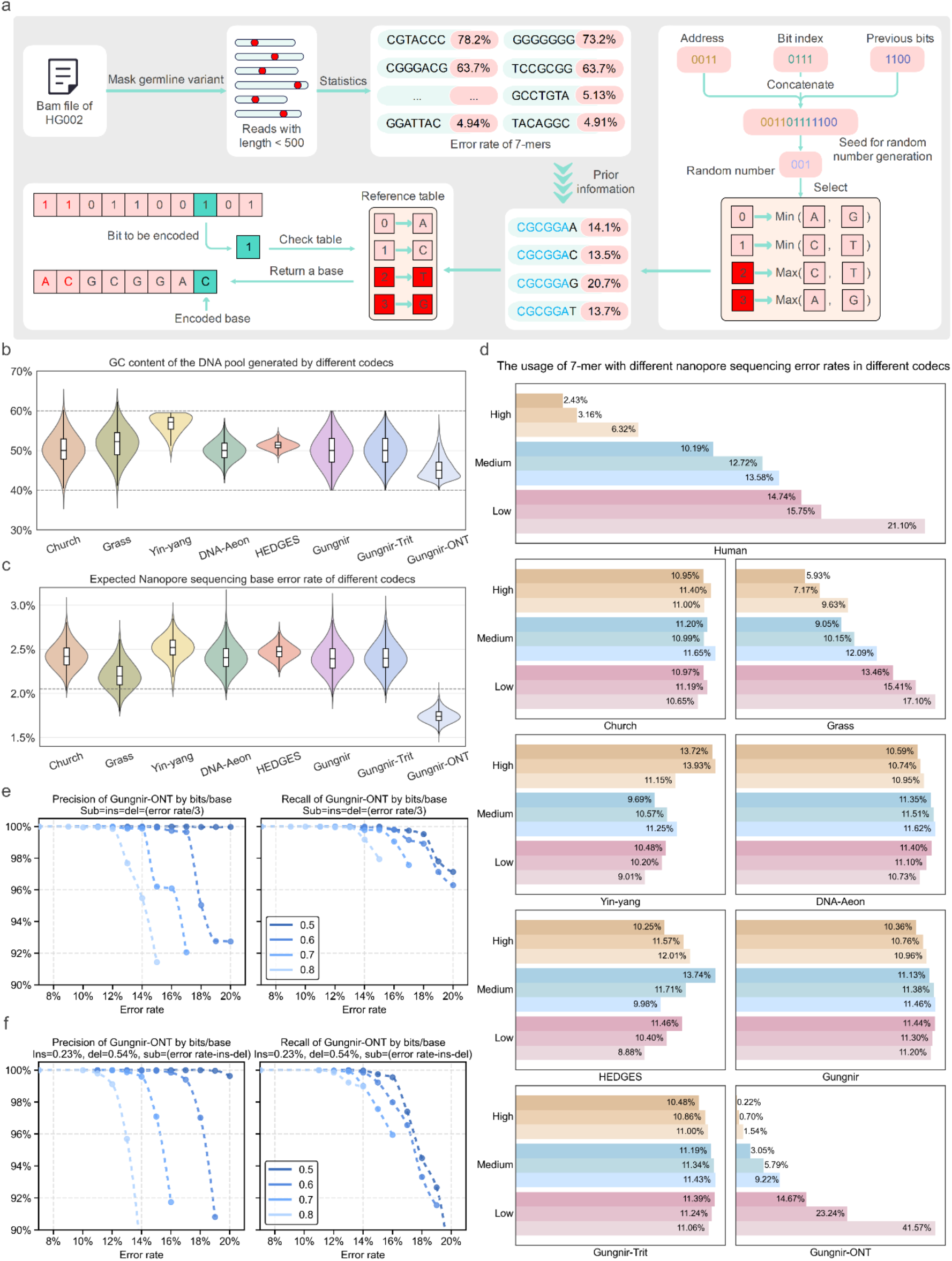
Details of Gungnir-ONT. a. A workflow of Gungnir-ONT. The principle of Gungnir-ONT is that during encoding, when multiple nucleotides can be used as the next base, the one that leads to a lower 7-mer sequencing error rate is chosen. b. The GC content of the DNA pool generated by different codecs. The dash lines at 40% and 60% mark an optimal range of GC content for Nanopore sequencing. c. Expected Nanopore sequencing base error rate of different codecs. The dash line marks the base error rate (∼2.08%) of Nanopore reads <500bp, using the standard sample HG002 as the example. d. The usage of 7-mers with known high, medium, and low Nanopore sequencing error rates in different codecs. The same in human genome is also provided for comparison. All 7-mers are sorted by known Nanopore sequencing error rate and divided into nine intervals with equal sizes. e-f.Precision and recall of Gungnir-ONT. *W*_*max*_= 20M was set. The data points in e-f are given in **Supplementary Table 4**.

**Figure 6c** illustrates the estimated Nanopore sequencing error rates for the generated DNA pools based on different codecs. Compared to the Gungnir, Gungnir-ONT significantly reduced the sequencing error rate by 28.4%. In comparison with other benchmarked methods, Gungnir-ONT significantly reduced errors by 19.6% to 29.9%. Notably, the maximum expected error rate for Gungnir-ONT was 2.08%, which is close to the observed Nanopore sequencing error rate of ∼2% for R10 chemistry. Furthermore, as depicted in **Figure 6d**, the 7-mer distribution of Gungnir-ONT closely resembled the natural human sequences, whereas the distributions produced by other benchmarked codecs exhibited a deviation. These results demonstrate Gungnir-ONT’s ability to effectively manage Nanopore sequencing errors.

**Figure 6e and 6f** depict the precision and recall of Gungnir-ONT, and the corresponding data recovery performance is presented in **Supplementary Figure 2**. For a fair comparison, we used the same setup described in the section **‘Flexible Information Density Options’**. Compared to Gungnir, Gungnir-ONT exhibited a reduction in error tolerance capacity by only 15% and 10% in in error profile 1 and 2, respectively. This reduction is attributed to the base preference introduced by the encoder, which diminishes the decoder’s ability to effectively prune erroneous branches when decoding sequences generated by Gungnir-ONT. Despite this slight loss of error tolerance, Gungnir-ONT remains crucial in frequent data retrieval scenarios due to its remarkable capability to reduce Nanopore sequencing errors.

## Discussion

DNA-based next-generation storage systems offer a radical departure from traditional silicon-based architectures ^23^, utilizing the unique properties of DNA molecules to store data to achieve a physical density of ∼200 petabytes per gram ^16^. In this paper, we developed Gungnir, a codec system that introduces a blockchain-inspired cryptographic framework to address the inherent limitations of conventional DNA storage codecs in terms of Indel correction. By reformulating sequence recovery as a proof-of-work problem, our system iteratively generates candidate sequences against cryptographic hashes, ensuring accurate reconstruction even at 20% error rates—a 4x improvement over existing methods. Our approach uniquely addresses substitutions and Indels in a sequence without resorting to excessive redundancy, achieving full data recovery while using 53% fewer DNA bases.

In our benchmark results, Gungnir showcases exceptional error tolerance, greatly exceeding the upper limit of error rates commonly encountered during DNA synthesis and sequencing. This remarkable feature ensures a substantial margin against errors introduced by DNA degradation during long-term storage, and facilitates data recovery from single reads. Unlike many other DNA codec systems ^11, 12, 13, 14, 15, 16^, Gungnir treats sequence addresses as unknown information to be guessed rather than explicitly encoded data. This design substantially improves the information density by saving 10–20 bits of address overhead.

These advancements enable Gungnir to maintain a balance between information density and error tolerance.

While conventional error correction methods suggest that substitutions are easier to handle than Indels, Gungnir achieves superior performance for sequences with a higher prevalence of Indels. This outcome is sensible: unlike substitutions and deletions, insertions do not result in the loss of any base information—they simply introduce additional bases, which are more manageable for our decoder. This finding offers a novel perspective that positions DNA-based codecs as a distinct issue instead of merely a derivative of computer error correction codes.

While our benchmarking focused on error rates lower than 20%, Gungnir’s architecture theoretically supports recovery from arbitrarily high error rates. However, higher error rates necessitate greater computational power due to the exponential increase in possible branches within the decoding algorithm. Despite this, advancements in computational power are likely to surpass DNA degradation rates, prompting us to investigate improved algorithms to further reduce computational complexity.

## Methods

### Overview of Gungnir

Gungnir is a DNA-based storage codec system that relies on substantial computing power for low-redundancy DNA storage. **Figure 1a** shows an overview of the Gungnir codec. The Gungnir codec system supports any type of information (such as images, videos, or audio) to be segmented into short binary fragments and subsequently transcoded into DNA sequences using the Gungnir encoder. The Gungnir encoder takes a binary fragment along with its corresponding address (i.e., the exact location of this fragment in the original information) as input and outputs an encoded sequence for DNA synthesis. The Gungnir decoder is designed for reconstructing the binary fragments from sequences with up to 20% of errors either from DNA synthesis, degradation during storage, or sequencing. Once all binary fragments are recovered, the original information can be accurately assembled according to the address in the fragment.

The error patterns introduced during DNA synthesis, degradation, and sequencing are different, platform-specific, and usually known in prior. To demonstrate that Gungnir can optimize for platform-specific error patterns, we implemented Gungnir-ONT, an extension of Gungnir that is optimized for Nanopore sequencing technologies with specific sequencing error patterns.

To demonstrate the versatility of Gungnir on a different balance between information density and tolerable error rate, we implemented Gungnir-Trit, a reconfigured Gungnir that has a higher information density (1.35 bits/base vs. 0.8 bits/base) while maintaining a moderate tolerable error rate (5% vs. 14%).

### Gungnir Transcoding

As depicted in **Figure 1b**, the Gungnir encoder concatenates the hash signature and data bits (payload) of a fragment into a binary stream and sequentially maps the bits into nucleotides. At each position, two of the A/C/G/T nucleotides are assigned to represent 0 and 1, while the other two nucleotides are utilized for redundancy in error correction. The specific mapping rules between bits (or trits) and bases for Gungnir, Gungnir-Trit, and Gungnir-ONT are discussed in following sections.

### Character Generation Rules for Gungnir

The Gungnir encoder generates a new reference table between the binary domain (0, 1) and the nucleotide alphabet *(A, C, T, G)* for each incoming bit, according to the previously encoded bases. As illustrated in **Figure 1b**, the encoding process begins by concatenating the 16-bit fragment address (i.e., the exact location of this fragment in original information), the 8-bit bit index (denoting the position of the target bit within the fragment), and the 40-bit history of previously encoded bits into a 64-bit transient seed. This seed serves as the input to a pseudorandom number generator for generating a 2-bit random number *r*. Within the finite field mod 4, the cyclic permutation (*r, r* + 1, *r* + 2, *r* + 3), defines an ordered mapping between the numerical sequence (0, 1, 2, 3) and the nucleotide (*A, C, T, G*). For example, when *r* = 2, the resulting permutation (2, 3, 0, 1) yields the correspondence (0 → *T*, 1 →*G*, 2 → *A*, 3 → *C*). This permutation (i.e., the reference table in **Figure 1b**) determines the encoding scheme: nucleotides *T* and *G* encode the information bits 0 and 1, respectively, while *A* and *C* are reserved as redundant bits for error correction purposes.

### Character Generation Rules for Gungnir-ONT

In Gungnir, the reference tables between the nucleotides *(A, C, T, G)* and digits (0, 1) are generated based on contextual information. While in Gungnir-ONT, base preferences specific to Nanopore sequencing technology are incorporated into the generated sequences. As depicted in **Figure 6a**, the prior information (with more details in the **Methods** section **‘Generation of Base Preference Table’**) is derived from the error rates observed in 7-mers distribution from a GIAB HG002 sample ^24^. Similar to the Gungnir, a 64-bit seed is generated based on the previously encoded bases. This seed then generates a random number *r* within the range [0,5], which partitions the set *(A, C, T, G)* into [(*C*_1_, *C*_2_); (*C*_3_, *C*_4_)]. The details of the partitions associated with different random number *r* are presented in **Supplementary Table 7**. Based on the partition, the bit message 0 is mapped to the character with lower error rate among (*C*_1_, *C*_2_), while the bit message 1 is mapped to the character with lower error rate among (*C*_3_, *C*_4_). For instance, as shown in **Figure 6a**, when *r* = 1, the partition of *(A, C, T, G)* results in [(*A, G*); (*C, T*)]. Since *A* and *C* exhibit lower error rates, the reference table is established as (0 → *A*, 1 → *C*).

### Character Generation Rules for Gungnir-Trit

In Gungnir and Gungnir-ONT, each base represents one bit, resulting in a maximum theoretical information density of 1 bit per base. For Gungnir-Trit, each base can accommodate one ternary digit (trit) rather than a bit. For a sequence of length 100, a maximum of 157 bits of information can be encoded. The transcoding rules of Gungnir-Trit are illustrated in **Supplementary Figure 3**. Following the concatenation of the payload and hash signature into a binary stream by the encoder, this binary stream undergoes conversion into its ternary representation (as detailed in **Supplementary Text 1**). Following the transcoding rules of Gungnir, a 101-bit seed is obtained for random number generation, comprising a 16-bit address, an 8-bit bit index, and 77 bits from the previous trits. The seed then generates a 2-bit random number *r*, with which a reference table between the trits (0, 1, 2) to nucleotide bases *(A, C, T, G)* is selected according to the same principles outlined in the **Methods** section **‘Character Generation Rules for Gungnir’**. In this configuration, the trits (0, 1, 2) represent valid bases, while only the base corresponding to 3 is rejected as redundancy for error correction.

### Biomedical Constraints-aware Transcoding

In our designed transcoding system, we proposed biomedical constraints-aware transcoding to adhere to several general biomedical constraints that could potentially limit DNA synthesis and sequencing. These constraints can be broadly categorized into three groups ^25^: GC content, homopolymers, and undesired motifs. To comply with these constraints, certain characters may be disabled at specific times, necessitating corresponding modifications in the transcoding rules. In this section, we will provide a detailed description of these constraints along with the associated transcoding rules.

#### GC Content

DNA-based data storage faces a critical constraint: maintaining the GC content of synthesized sequences within the optimal range of 40% to 60%. An imbalanced GC content can lead to secondary structure formation ^25^ or non-uniform sequence coverage ^26^ during synthesis and sequencing. To address this, the Gungnir encoder incorporates a real-time GC content balancing mechanism. If the GC content exceeds the specified range, the encoder dynamically adjusts nucleotide selection by excluding non-compliant bases. For instance, consider a reference table where (0 → *G*, 1 → *A*, 2 → *C*, 3 → *T*), if the GC content among previously encoded bases exceeds 60%, the *G* and *C* should be disabled, resulting in a new reference table of (0 → *A*, 1 → *T*).

#### Homopolymers

Homopolymers, defined as continuous repeats of identical nucleotides, pose great challenges for accurate sequencing. Nanopore sequencing methods typically exhibit reduced performance in determining the lengths of homopolymer ^27^, especially when they exceed 5 bases ^28^. To address this issue, the Gungnir encoder is designed to prevent homopolymers longer than 3 bases by excluding corresponding characters. For instance, if the original reference table in a position is (0 → *T*, 1 → *G*, 2 → *A*, 3 → *C*), but the previous 3 characters are “*GGG*”, the *G* should be disabled to prevent a homopolymer of length 4.

Consequently, the new reference table becomes (0 → *T*, 1 → *A*).

#### Undesired Motifs

Biologically functional motifs are crucial sequence elements that necessitate systematic consideration. For instance, restriction enzyme recognition sites ^29^ are essential for initiating the amplification steps of various workflows. Consequently, due to the limitations associated with synthesis, it is crucial to exclude such motifs during the encoding process. Gungnir implements this functionality as optional, with the encoder maintaining a predefined library of undesired motifs. For instance, 6-mer “*GTGCAC*” is a restriction enzyme recognition in this library. When the previous 5-mer “*GTGCA*” is detected, the *C* should be disabled at the next position to prevent the occurrence of this motif.

### Generation of Base Preference Table

The base preference table is an optional module within the Gungnir codec system, which is designed to assign scores to each k-mer. K-mers with higher scores are more likely to be selected by the Gungnir encoder. This table can be informed by relevant prior knowledge, such as Nanopore sequencing error rates, enabling the generation of sequences that are preferred in specific target scenarios.

Gungnir-ONT is an implementation that utilizes a base preference table to minimize sequencing errors. The value of *k* is set to 7 to align with the characteristics of Nanopore sequencing. To generate the base preference table, we reviewed all reads in a 50-fold GIAB HG002 sample^24^ and filtered out reads longer than 500 bases. Using the ONT variant caller Clair3 ^30^, we masked undesired germline variants and counted the occurrences of each 7-mer. Substitutions, deletions, and insertions at different positions within each 7-mer were recorded separately and summarized to obtain a total error rate. Subsequently, the Gungnir-ONT encoder can assign scores to different bases based on these error rates.

### Decoding Algorithm of Gungnir

The Gungnir decoder employs a proof-of-work framework that combines exhaustive search with cryptographic hash signature validation to recover the data bits from a single sequence copy containing erroneous bases. Unlike the conventional one-pass decoding strategies, Gungnir decoder adopts a progressive verification mechanism inspired by blockchain architectures. The framework employs a hash signature validation scheme based on murmur3 ^31^, a renowned hash function celebrated for its exceptional speed and randomness, to provide robust mistake detection capabilities. Within a proof-of-work-style decoding algorithm, the decoder systematically explores the solution space until it identifies a candidate that satisfies all cryptographic validation criteria. This section will delve into the technical details of the Gungnir decoder.

### Generation of Initial Hypothesis

As depicted in **Figure 2a**, the Gungnir decoder is a variant of the A* algorithm ^32^ to explore potential hash matches through a dynamically constructed hypothesis tree (**Figure 2b**). In each hypothesis, Gungnir guesses the address and the binary stream simultaneously, and tracks the corresponding positions in the sequence copy. Additionally, the hypothesis accumulates an edit distance to prune the incorrect branches.

When decoding, the decoder generates initial hypotheses without considering prior knowledge of the exact location (address) of the data bits (payload). Each address is assigned with an initial hypothesis, assuming an empty binary stream and zero edit distance. However, for branches stemming from initial hypotheses with an incorrect address, as detailed in the section **‘Character Generation Rules for Gungnir’**, the downstream seed for random number generation will invariably be incorrect. This results in an erroneous reference table between (0, 1) and (*A, C, T, G*). Consequently, hypotheses with an incorrect address will be pruned in a very early stage due to their higher accumulated edit distance (details in the **Methods** section **‘Construction of the Hypothesis Tree’**) or fail hash validation (details in the **Methods** section **‘Hash Signature Validation in a Hypothesis’**).

### Generation of Children Hypothesis from a Parent Hypothesis

**Figure 2c** illustrates how a parent hypothesis generates its children hypotheses. Following the transcoding rules, a new reference table between (0, 1) and *(A, C, T, G)* at the new position can be selected based on the guessed information within the parent hypothesis. For instance, if the reference table is selected as (0 − *C*_0_, 1 − *C*_1_), then either C_0_ or C_1_ will be permitted in the subsequent position. In the sequence to be decoded, the base to be looking at is designated as C_N_. To ascertain what happened to C_N_, five children hypotheses are generated:

1. Match or Substitution (two children hypotheses). In this case, the decoder guesses that the C_N_ is either C_0_ or C_1_, corresponding to the digit 0 or 1. If the guessed base (C_0_ or C_1_) differs from C_N_, the decoder assumes a substitution error here and accumulates the edit distance by 1. For example, in **Figure 2b**, if the decoder guessed that the next bit is 0, the C_0_ will be G. However, in the sequence to be decoded, C_N_ is C, resulting in an assumption of substitution of G→C. Similarly, if the decoder guessed that the next bit is 1, then C_1_ matches C_N_, so no edit distance is accumulated.
2. Deletion (two children hypotheses). In this case, the decoder guesses that a deletion has occurred prior to C_N_. Consequently, the decoder disregards C_N_ and attempts to guess the deleted base to be either C_0_ or C_1_, with edit distance accumulated by 1 for both children hypotheses.
3. Insertion (one children hypothesis). In this case, the decoder guesses that an insertion has occurred at C_N_, indicating that C_N_ is not a base in the original sequence. Consequently, the decoder refrains from attempting to guess any new bit and accumulates the edit distance by 1.

### Construction of the Hypothesis Tree

In an initial hypothesis, all bits remain unguessed, and the edit distance is set to zero. Following the children hypothesis generation rules, bits are guessed in the generation of subsequent hypotheses. As illustrated in **Figure 2a**, the initial layer comprises hypotheses corresponding to all possible addresses. Then each hypothesis within a layer generates five sub-hypotheses (as detailed in the **Methods** section **‘Generation of Children Hypothesis from a Parent Hypothesis’**), and accumulating edit distance for any substitutions, deletions, or insertions. To maintain computational tractability, the decoder implements a constraint *W*_*max*_ (maximum hypotheses allowed): when the number of hypotheses exceeds *W*_*max*_, the candidates with highest edit distance are pruned. Once a hypothesis has completed guessing, the leaf is retained in the following steps and stops children generation. The hypothesis tree construction stops only when all preserved hypotheses completed guessing. According to the transcoding rules, if a parent hypothesis is incorrect regarding the address, previous bits, or bit index, the reference table for its children hypotheses is also incorrect.

This leads to a substantial increase in the edit distance of children hypotheses, ultimately resulting in hypotheses pruning. Hence, the remaining leaf hypothesis is more likely to be correct for subsequent hash signature validation.

### Hash Signature Validation in a Hypothesis

Upon completing the construction of the hypothesis tree, the decoder generates a set of hypotheses, each containing the exact expected number of bits for hash signature validation. For clarity, *H* represents the hash signature, *A* denotes the address, and *P* signifies the payload. As depicted in **Figure 2a**, the decoder validates each hypothesis by verifying the equation *hash*(*A, P*) = *H*. Hypotheses that meet this condition are retained, while those that fail verification are discarded. The cryptographic properties of the murmur3 hash function ensure that even minor errors in *A* or *P* result in a drastically different *H*, thereby safeguarding the validation process and retaining only the successful hypotheses. Following this verification, successful hypotheses are added to a candidate set, from which the hypothesis with the minimal edit distance is selected as the decoding result.

However, if the *W*_*max*_ (maximum hypotheses allowed) is too small, the correct branch may be pruned, potentially leaving an empty candidate set. In such instances, the decoder would increase the *W*_*max*_ and try the decoding algorithm again until a valid output is obtained.

Theoretically, with a sufficiently large *W*_*max*_, the correct branch will ultimately be included in the candidate set.

### An Advancing Cutoff for Edit Distance

The Gungnir decoder employs an advancing cutoff for edit distance (*ED*_*max*_) as an optional parameter to control decoding efficiency. Initially, it is set to a small value, such as 3. During decoding, the decoder prunes the hypothesis with an accumulated edit distance exceeding *ED*_*max*_. When decoding, *W*_*max*_ is increased for sequences with decoding failures at this *ED*_*max*_ to provide more computational power. If failures persist due to sequences containing more errors than *ED*_*max*_ allows, the decoder increases *ED*_*max*_ and restarts the decoding algorithm. This process iterates until all sequences are decoded or reaching upper limit of *ED*_*max*_, such as *ED*_*max*_ = 20 in a sequence of length 100.

### Configurations for Different Information Density

In each sequence, the information density is defined as 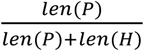 bits/base, where *P* represents the payload and *H* denotes the hash signature. A longer *H* reduces the likelihood of hash collisions, allowing the decoder to identify the correct hypothesis more accurately from the candidate set after hash signature validation. However, when the total sequence length remains constant, increasing the hash signature length results in a corresponding decrease in payload length, leading to a reduction in information density. Consequently, the configurations of Gungnir involves a trade-off between information density and error tolerance capacity. To accommodate the diverse requirements of different scenarios, we provide various configurations, spanning from 0.5 to 1.45 bits/base. In this section, we will delve into these configurations.

### Default Configuration: Better Information Density

The default configuration is effective when the length of the hash signature is relatively short (information density ≥0.8 bits/base for Gungnir and Gungnir-ONT, information density ≥ 1.35 bits/base for Gungnir-Trit). As illustrated in **Supplementary Figure 4a**, the binary stream can be represented as [*H*; *P*], and the hash validation between *H* and *hash*(*A, P*) will be performed after all digits have been guessed.

### Multiple Hash Validation Configuration: Better Error Tolerance

Our default configuration operates effectively for Gungnir when error rates remain below 10%. However, at elevated error rates exceeding 15%, the primary challenge of the decoding algorithm shifts from the reliability of hash signature validation to preventing the decoder from pruning correct branches. Specifically, correct branches may accumulate higher edit distances and are more likely to be incorrectly pruned, especially in regions of the sequence to be decoded characterized by a high concentration of errors. Consequently, increasing the length of *H* may not improve the decoding success rate. To address this, multiple hash validation proves to be more effective by frequently pruning incorrect branches during hypothesis tree construction. This configuration is applied when the length of the hash signature is relatively long (information density <0.8 bits/base for Gungnir and Gungnir-ONT, information density <1.35 bits/base for Gungnir-Trit).

In this configuration, the digits of *P* are divided into a set (*p*_0_, *p*_1_, …, *p*_*k*_), where the length of each *p*_*i*_ is approximately equal. As illustrated in **Supplementary Figure 4b**, the hash signature *H* is not a single number but rather a set (*h*_0_, *h*_1_, …, *h*_*k*_). The binary stream can be represented as [*h*_0_; *h*_1_; *p*_1_; *h*_2_; *p*_2_, …, *h*_*k*_; *p*_*k*_; *p*_0_]. Within this information stream, *h*_0_ serves as the overall hash signature, computed as *h*_0_ = *hash*(*A, p*_0_, *p*_1_, …, *p*_*k*_), or equivalently, *h*_0_ = *hash*(*A, P*). For *h*_*i*_ (1 ⩽ *i* ⩽ *k*), is calculated as *h*_*i*_ = *hash*(*A, h*_0_, *p*_*i*_). During the decoding process, a hash validation is performed for each *h*_*i*_ (0 ⩽ *i* ⩽ *k*). The overall hash signature validation for *h*_0_ is conducted after the construction of the hypothesis tree has been completed (as detailed in the **Methods** section **‘Construction of the Hypothesis Tree’**).

For any other *h*_*i*_ (1 ⩽ *i* ⩽ *k*), hash validation occurs once expected number of digits in a hypothesis have been guessed, and hypotheses that fail any hash validation are immediately pruned. For example, once all digits of *h*_0_, *h*_1_ and *p*_1_ have been guessed in a hypothesis, the decoder verifies the equality *h*_1_ = *hash*(*A, h*_0_, *p*_1_), promptly pruning incorrect branches if a hypothesis fails this validation. By doing this, correct hypothesis can be preserved when the decoder navigates sequence regions of high error concentration.

## Acknowledgments

R.L. was supported CRF (C7003-24Y) of the Research Grants Council (RGC) of Hong Kong, and the URC fund at HKU.

## Author contributions

Conceptualization: J. Z., R. L. Methodology: J. Z., R. L. Investigation: J. Z. Software: J. Z., Z.

Z. Visualization: J. Z., J. S. Validation: J. S., J. Z. Supervision: R. L. Writing—original draft: J. Z., Z. Z., J. S., R. L. Writing—review & editing: All authors.

## Code availability

Gungnir is available open-source at https://github.com/HKU-BAL/Gungnir under the BSD 3-Clause license.

## Data availability

The authors declare that all data supporting the findings, including source data and analysis results of this study are available at http://www.bio8.cs.hku.hk/gungnir/.

