## Supplementary Materials for "Guess till correct: Gungnir codec enabling high error-tolerance and low-redundancy DNA storage through substantial computing power"

### Supplementary Figures

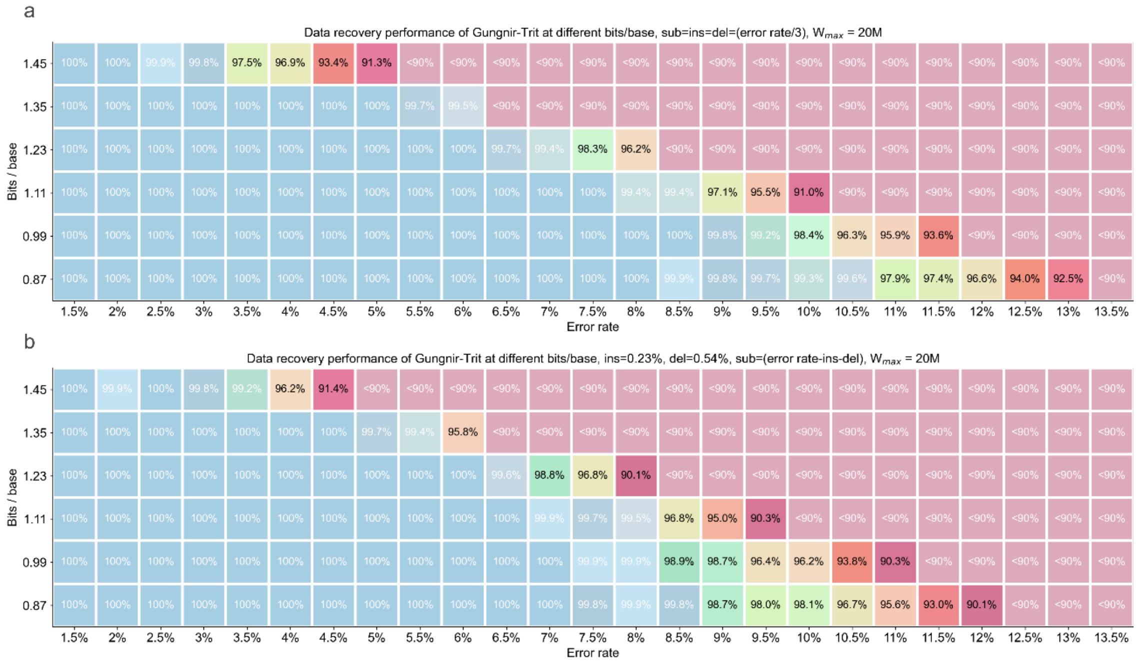

#### Supplementary Figure 1. Data recovery performance of Gungnir-Trit.

Data recovery performance of Gungnir-Trit at different bits/base and two different error profiles with constrained memory ($W_{max}$ (maximum hypotheses allowed) =20M, ~20GB). a. Percentage of data recovered by Gungnir-Trit in error profile 1 (equal proportions of substitutions, insertions, and deletions errors). b. Percentage of data recovered by Gungnir-Trit in error profile 2 (0.54% deletions and 0.23% insertions, gradually increasing substitutions).

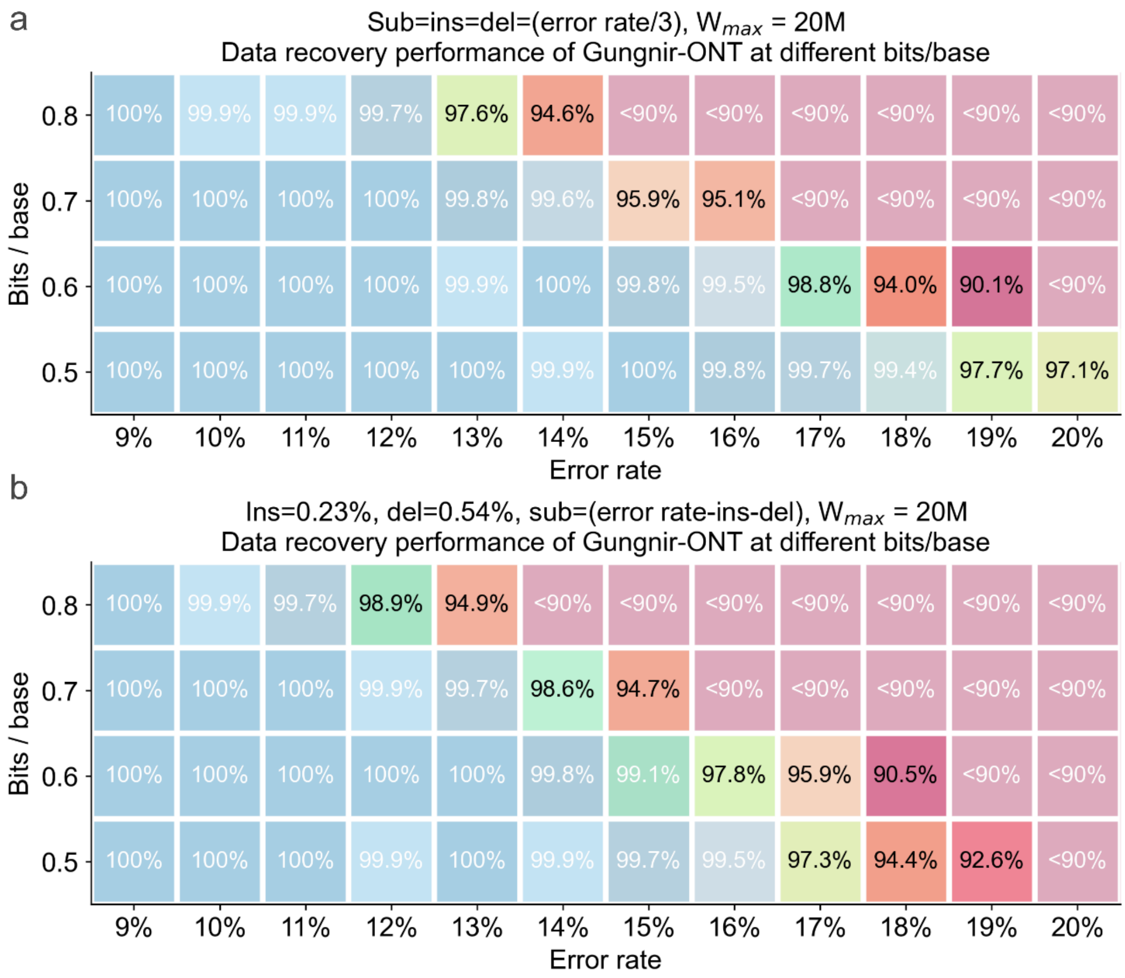

### Supplementary Figure 2. Data recovery performance of Gungnir-ONT.

Data recovery performance of Gungnir-ONT at different bits/base and two different error profiles with constrained memory ($W_{max}$ (maximum hypotheses allowed) =20M, ~20GB). a. Percentage of data recovered by Gungnir-ONT in error profile 1 (equal proportions of substitutions, insertions, and deletions errors). b. Percentage of data recovered by Gungnir-ONT in error profile 2 (0.54% deletions and 0.23% insertions, gradually increasing substitutions).

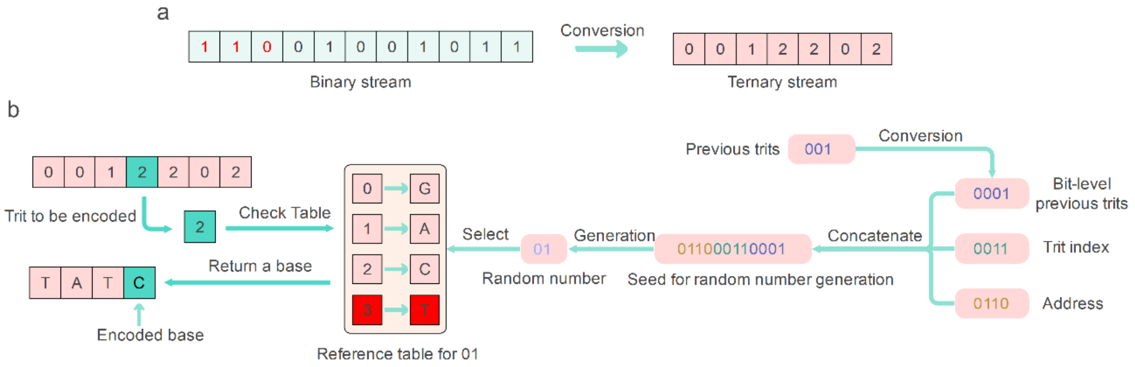

### Supplementary Figure 3. Character generation rule for Gungnir-Trit.

a. After the encoder concatenates the message bits (payload, labeled in black) and the hash signature (labeled in red) into a binary stream, this stream is converted into a ternary representation, with every 7 ternary digits (trit) represent 11 bits. b. Generation of a new base. The encoder transcodes trits into nucleotides trit-by-trit, with which nucleotide to be used for a trit decided by one of the preset reference tables, chosen by a random number generated from a seed constructed as the concatenation of address, trit index, and previous trits.

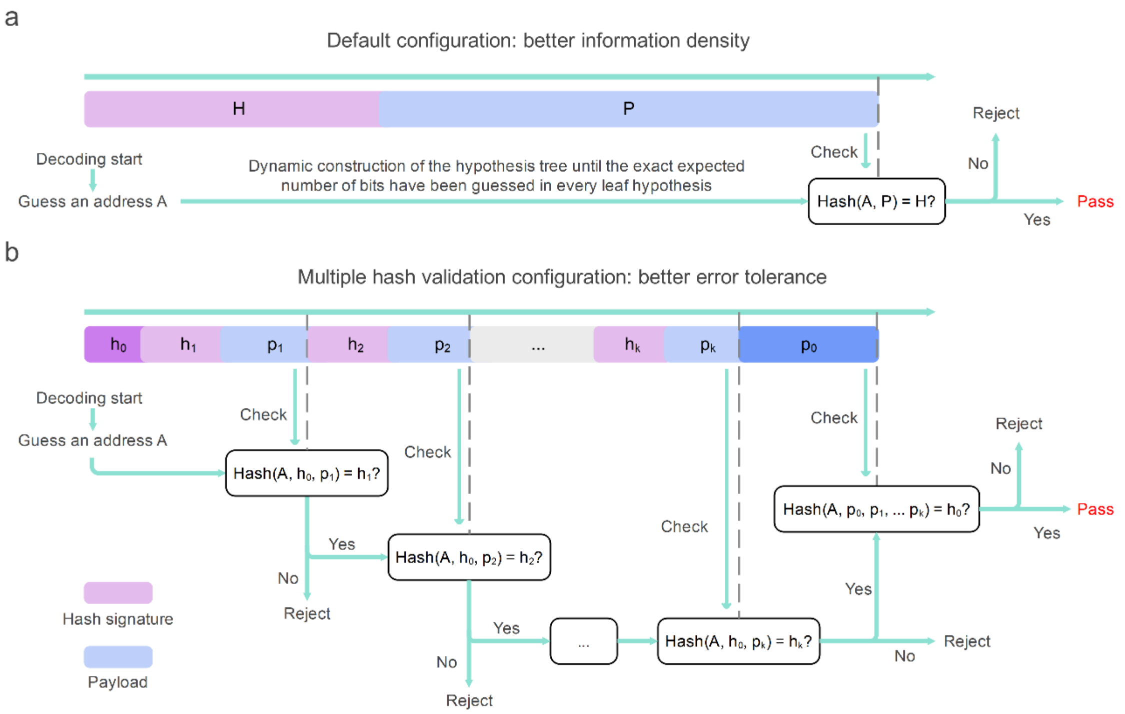

### Supplementary Figure 4. Configurations for different bits/base.

a. An overview of the default configuration (information density $\geqslant$0.8 bits/base for Gungnir and Gungnir-ONT, information density $\geqslant$1.35 bits/base for Gungnir-Trit). The information stream is the concatenation of the payload ($P$) and the digital signature ($H$) of a fragment, and the address ($A$) is utilized as a random seed. Here, $H$ is derived from $hash(A, P)$, and the information stream can be represented as $[H; P]$. During decoding, the hash validation is performed after the exact expected number of digits have been guessed. b. An overview of the multiple hash validation configuration (information density <0.8 bits/base for Gungnir and Gungnir-ONT, information density <1.35 bits/base for Gungnir-Trit). Here the full payload $P$ is divided into $(p_0, p_1, \dots, p_k)$, while the hash signature $H$ is not a single number but rather a set $(h_0, h_1, \dots, h_k)$. The information stream can be represented as $[h_0; h_1; p_1; h_2; p_2, \dots, h_k; p_k; p_0]$. During decoding, the hash validation is performed after the exact expected number of digits in each $p_i$ have been guessed.

### Supplementary Tables

#### Supplementary Table 1. Data recovery performance of different codecs.

This table shows the details of data recovery performance across various codec systems (**Figure 3a**). Here the sequence length of Gungnir is fixed at 100. No upper limit of $W_{max}$ (maximum hypotheses allowed) has been set in Gungnir decoder. Two different error profiles were tested (error profile 1: sub=ins=del=(error rate/3); error profile 2: ins=0.23%, del=0.54%, sub=(error rate-ins-del)). Sub, ins, and del stand for substitution, insertion, and deletion, respectively. NA indicates that the error rate was not applied in the error profile.

| Church | | | | | | | | | |
|---|---|---|---|---|---|---|---|---|---|
| Error rate | 0.3% | 0.5% | 0.75% | 0.77% | 1% | 1.5% | 2% | 2.5% | 3% |
| Error profile 1 | 71.6% | 58.9% | 45.0% | NA | 35.3% | 20.8% | 12.7% | 6.9% | 4.4% |
| Error profile 2 | NA | NA | NA | 44.3% | 32.7% | 18.3% | 10.6% | 6.1% | 3.1% |
| **Grass** | | | | | | | | | |
| Error rate | 0.3% | 0.5% | 0.75% | 0.77% | 1% | 1.5% | 2% | 2.5% | 3% |
| Error profile 1 | 75.5% | 62.6% | 49.9% | NA | 40.1% | 28.3% | 15.6% | 9.6% | 5.8% |
| Error profile 2 | NA | NA | NA | 41.9% | 36.1% | 30.1% | 20.8% | 14.8% | 10.9% |
| **Yin-yang** | | | | | | | | | |
| Error rate | 0.3% | 0.5% | 0.75% | 0.77% | 1% | 1.5% | 2% | 2.5% | 3% |
| Error profile 1 | 81.5% | 73.0% | 61.0% | NA | 48.0% | 33.3% | 21.2% | 14.3% | 9.2% |
| Error profile 2 | NA | NA | NA | 44.3% | 39.2% | 31.7% | 22.5% | 16.1% | 10.6% |

| DNA-Aeon | | | | |
|---|---|---|---|---|
| Error rate | 3.6% | 3.9% | 4.2% | 4.5% |
| Error profile 1 | 100% | 100% | 0% | 0% |
| Error profile 2 | 100% | 100% | 100% | 0% |

| HEDGES | | | | | |
|---|---|---|---|---|---|
| Error rate | 4.5% | 5.25% | 6% | 6.75% | 7.5% |
| Error profile 1 | 100% | 100% | 90% | 30% | 0% |
| Error profile 2 | 100% | 100% | 95% | 10% | 0% |

| Gungnir (0.5 bits/base) | | | | | | |
|---|---|---|---|---|---|---|
| Error rate | 10% | 12% | 14% | 16% | 18% | 20% |
| Error profile 1 | 100% | 100% | 100% | 100% | 100% | 100% |
| Error profile 2 | 100% | 100% | 100% | 100% | 100% | 100% |

### Supplementary Table 2. The maximum tolerable error rate at different bits/base of different codecs.

This table shows the comparison of the maximum tolerable error rate at different bits/base of different codecs (**Figure 3b**). Here the sequence length of Gungnir is fixed at 100. No upper limit of $W_{max}$ (maximum hypotheses allowed) has been set in Gungnir decoder. Two different error profiles were tested (error profile 1: sub=ins=del=(error rate/3); error profile 2: ins=0.23%, del=0.54%, sub=(error rate-ins-del)). Sub, ins, and del stand for substitution, insertion, and deletion, respectively. A codec system is considered capable of tolerating an error rate only if all payloads are successfully recovered. Conversely, NA indicates that a codec system cannot handle the minimum error rate introduced in an error profile presented in **Supplementary Table 1**.

| Codec system | Information density (bits/base) | Error profile 1 | Error profile 2 |
|---|---|---|---|
| Church | 0.83 | NA | NA |
| Yin-yang | 0.50 | NA | NA |
| DNA-Aeon | 0.73 | 3.90% | 4.02% |
| Hedges | 0.64 | 5.25% | 5.25% |
| Gungnir | 0.5 | 20% | 20% |
| | 0.6 | 18% | 17% |
| | 0.8 | 14% | 12% |
| Gungnir-ONT | 1.11 | 8% | 7.5% |
| | 1.35 | 5% | 4.5% |
| | 1.45 | 2% | 1.5% |

### Supplementary Table 3. Computational cost analysis of Gungnir.

The table shows the details of the count of sequences that can be successfully decoded with various $W_{max}$ (maximum hypotheses allowed) and the corresponding memory consumption shown in **Figure 4a and 4b**. Here the configuration of Gungnir is 0.5 bits/base, and the sequence length is fixed at 100. Error profile 1 is tested, where the number of substitutions, insertions, and deletions is equal.

| Rounds | Error rate | Round 1 | Round 2 | Round 3 | Round 4 | Round 5 |
|---|---|---|---|---|---|---|
| Number of sequences successfully decoded | 10% | 3,108 | 0 | 0 | 0 | 0 |
| | 12% | 3,100 | 8 | 0 | 0 | 0 |
| | 14% | 2,997 | 102 | 8 | 1 | 0 |
| | 16% | 2,863 | 196 | 42 | 7 | 0 |
| | 18% | 2,584 | 422 | 78 | 21 | 3 |
| | 20% | 2,271 | 562 | 167 | 87 | 21 |
| Time (minutes) | 10% | 0.57 | 0 | 0 | 0 | 0 |
| | 12% | 0.54 | 0.07 | 0 | 0 | 0 |
| | 14% | 0.77 | 0.23 | 0.12 | 0.11 | 0 |
| | 16% | 1.08 | 0.49 | 0.35 | 0.29 | 0 |
| | 18% | 1.26 | 1.32 | 0.87 | 1.21 | 1.28 |
| | 20% | 1.20 | 2.21 | 2.43 | 7.98 | 11.59 |
| $W_{max}$ | | 100 K | 1 M | 5 M | 20 M | 200 M |
| Peak memory | | 100 MB | 1 GB | 5 Gb | 20 GB | 200 GB |

### Supplementary Table 4. Data recovery performance of Gungnir at different bits/base.

The table shows the details of precision, recall and data recovery performance of Gungnir, Gungnir-Trit and Gungnir-ONT. Here the sequence length is fixed at 100, both error profiles are tested. The $W_{max}$ (maximum hypotheses allowed) is set as 20M (~20GB memory). Error rate added at different bits/base will gradually increase, until it comes to 20% or data recovery is lower than 90%. Details of this table can be found in **Supplementary Table 4.xlsx**.

#### Supplementary Table 5. Precision and recall of Gungnir at different sequence lengths.

The table shows the details of the precision and recall of Gungnir at different length (**Figure 5a and 5b**). Here the configuration of Gungnir is 0.9 bits/base, and sequence length is set as 100, 200 and 300. Error profile 1 is tested, where the number of substitutions, insertions, and deletions is equal. The $W_{max}$ (maximum hypotheses allowed) is set as 100K (~100MB memory). Details of this table can be found in **Supplementary Table 5.xlsx**.

#### Supplementary Table 6. Effect of the gradually advancing edit-distance cutoff ($ED_{max}$) on decoding mistakes and decoding failures.

The table shows the details of the comparison of the number of decoding mistakes and decoding failures when a gradually advancing edit-distance cutoff ($ED_{max}$) is applied or not (**Figure 5c**). Here the configuration of Gungnir is 0.8 bits/base, and sequence length is fixed at 100. Error profile 1 is tested, where the number of substitutions, insertions, and deletions is equal. The $W_{max}$ (maximum hypotheses allowed) is set as 20M (~20GB memory).

| With/Without $ED_{max}$ | Error rate | Number of sequences | Number of decoding mistakes | Number of decoding failures |
|---|---|---|---|---|
| With $ED_{max}$ | 15% | 1943 | 1 | 0 |
| | 16% | 1943 | 16 | 5 |
| | 17% | 1943 | 31 | 19 |
| | 18% | 1943 | 258 | 8 |
| Without $ED_{max}$ | 15% | 1943 | 53 | 1 |
| | 16% | 1943 | 97 | 0 |
| | 17% | 1943 | 194 | 3 |
| | 18% | 1943 | 300 | 9 |

### Supplementary Table 7. Partitions of bases associated with different random number $r$ for Gungnir-ONT.

The table shows the details of the partitions of four bases A/C/G/T associated with different random number $r$ for Gungnir-ONT. Here $r$ is an integer between 0 and 5. For a certain $r$, 0 and 1 represent the base with minimal expected Nanopore sequencing error (valid bits), while 2 and 3 represent the base with maximum expected Nanopore sequencing error (redundancy for error correction).

| Bits | 0 | 1 | 2 | 3 |
|---|---|---|---|---|
| r=0 | (A, C) | (G, T) | (G, T) | (A, C) |
| r=1 | (A, G) | (C, T) | (C, T) | (A, G) |
| r=2 | (A, T) | (C, G) | (C, G) | (A, T) |
| r=3 | (C, G) | (A, T) | (A, T) | (C, G) |
| r=4 | (C, T) | (A, G) | (A, G) | (C, T) |
| r=5 | (G, T) | (A, C) | (A, C) | (G, T) |

### Supplementary Texts

#### Supplementary Text 1. Configurations for benchmarked DNA storage codecs.

##### Church

Church et al. [1] reports encoding 96 bits in each sequence of length 115, so the information density is $\frac{96}{115} \approx 0.83$ bits/base. In their strategy, the maximum length of homopolymers is 3, and no GC content constraints have been set. The code is sourced from the state-of-the-art simulator Chamaeleo [2].

##### Yin-yang

Ping et al. [3] reports encoding 256 bits in each sequence of length 160 in the Figure 3 of their paper, where 16 nucleotide for address and 144 nucleotide for a data package with Reed-Solomon code (n=18, k=16). Actually, in their GitHub repository https://github.com/ntpz870817/DNA-storage-YYC.git, the length of address is dynamic rather than an adjustable fixed parameter, while some additional binary streams will be generated for encoding. Following their 144-nucleotide-data-package design, we encoded the “The Ugly Duckling” (155,392 bits) into 667 sequences of length 155, so the actual information density is $\frac{155392}{667*155} \approx 1.50$ bits/base. Here the maximum length of homopolymers is set as 3, and the GC content is constrained within 40% and 60%.

##### Grass

Grass et al. [4] reports encoding 160 bits in each sequence of length 117, so the information density is $\frac{160}{117} \approx 1.37$ bits/base. In their strategy, the maximum length of homopolymers is 3, and no GC content constraints have been set. The code is sourced from the state-of-the-art simulator Chamaeleo.

##### DNA-Aeon

Welzel et al. [5] reports encoding 112 bits in each sequence of length 110, and using DNA-fountain [6] as outer code with 40% supplementary sequences generated, so the information density is $\frac{112}{110} \times \frac{1}{1+40\%} \approx 0.73$ bits/base. Following their default setting, the GC content is constrained between 40% and 60% in 10 bp intervals, and the maximum length of homopolymers is 3. The code is sourced from their GitHub repository https://github.com/MW55/DNA-Aeon.git.

##### Hedges

Press et al. [7] reports encoding a maximum of 254 bits in each sequence of length 300, and using Reed-Solomon code (n=255, k=223) as outer code for error correction. Actually, in their GitHub repository https://github.com/whpress/HEDGES.git, the defaulting setting encodes 216 bits in each sequence of length 294. Here we follow their GitHub repository, calculate the information density as $\frac{216}{294} \times \frac{223}{255} \approx 0.64$ bits/base. Following their default setting, the GC number is constrained between 4 and 8 in 12 bp intervals. For fair comparison, the maximum length of homopolymers is set as 3.

#### Gungnir

In all experiments, the GC content is constrained between 40% and 60%, and the maximum homopolymer run is set as 3, with no undesired motifs set. Some optional configurations, including 0.5, 0.6, 0.7 and 0.8 bits/base are offered for flexibility. The code of Gungnir can be found in the GitHub repository https://github.com/HKU-BAL/Gungnir.

#### Gungnir-Trit

The code of Gungnir-Trit can be found in the GitHub repository of Gungnir. Gungnir-Trit follows the same maximum homopolymer run, GC content and undesired motifs set of Gungnir, with flexible configurations including 0.87, 0.99, 1.11, 1.23, 1.35 and 1.45 bits/base. In Gungnir-Trit, the length of sequence is always set as 100 bases, corresponding to 100 trits. To facilitate the multiple hash validation in **Supplementary Figure 4b**, the 100 trits can represent 157 bits, with each 7 trits represent 11 bits, and the last 2 trits represent 3 bits. In this way, the hash validation can be performed after all 11 bits corresponding to a 7-trit segment have been guessed by the decoder.

#### Gungnir-ONT

The code of Gungnir-ONT can be found in the GitHub repository of Gungnir. Gungnir-ONT follows the same maximum homopolymer run, GC content, undesired motifs and information density set of Gungnir. The base preference table of Nanopore sequencing can be found in http://www.bio8.cs.hku.hk/gungnir/.
